# Genetic screen suggests an alternative mechanism for azide-mediated inhibition of SecA

**DOI:** 10.1101/173039

**Authors:** Rachael Chandler, Mohammed Jamshad, Jack Yule, Ashley Robinson, Farhana Alam, Karl A. Dunne, Naomi Nabi, Ian Henderson, Damon Huber

**Affiliations:** Institute for Microbiology and Infection, University of Birmingham, Birmingham, United Kingdom; Department of Cancer Sciences, University of Birmingham, Birmingham, UK; Department of Biological Sciences, University of Huddersfield, Huddersfield, UK

**Keywords:** Sec, SecA, azide, protein translocation, TraDIS, phospholipid binding, metal binding

## Abstract

Sodium azide prevents bacterial growth by inhibiting the activity of SecA, which is required for translocation of proteins across the cytoplasmic membrane. Azide inhibits ATP turnover *in vitro*, but its mechanism of action *in vivo* is unclear. To investigate how azide inhibits SecA in cells, we used transposon directed insertion-site sequencing (TraDIS) to screen a library of transposon insertion mutants for mutations that affect the susceptibility of *E. coli* to azide. Insertions disrupting components of the Sec machinery generally increased susceptibility to azide, but insertions truncating the C-terminal tail (CTT) of SecA decreased susceptibility of *E. coli* to azide. Treatment of cells with azide caused increased aggregation of the CTT, suggesting that azide disrupts its structure. Analysis of the metal-ion content of the CTT indicated that SecA binds to iron and the azide disrupts the interaction of the CTT with iron. Azide also disrupted binding of SecA to membrane phospholipids, as did alanine substitutions in the metal-coordinating amino acids. Furthermore, treating purified phospholipid-bound SecA with azide in the absence of added nucleotide disrupted binding of SecA to phospholipids. Our results suggest that azide does not inhibit SecA by inhibiting the rate of ATP turnover *in vivo*. Rather, azide inhibits SecA by causing it to “backtrack” from the ADP-bound to the ATP-bound conformation, which disrupts the interaction of SecA with the cytoplasmic membrane.

**Significance statement:** SecA is a bacterial ATPase that is required for the translocation of a subset of secreted proteins across the cytoplasmic membrane. Sodium azide is a well-known inhibitor of SecA, but its mechanism of action *in vivo* is poorly understood. To investigate this mechanism, we examined the effect of azide on the growth of a library of ∼1 million transposon insertion mutations. Our results suggest that azide causes SecA to backtrack in its ATPase cycle, which disrupts binding of SecA to the membrane and to its metal cofactor, which is iron. Our results provide insight into the molecular mechanism by which SecA drives protein translocation and how this essential biological process can be disrupted.

## Introduction

In bacteria, protein substrates of the Sec machinery are transported across, or inserted into, the cytoplasmic membrane through a channel composed of integral membrane proteins SecY, -E and -G (Sec61α, -β and –γ in eukaryotes). In bacteria, translocation of most soluble periplasmic proteins and outer membrane proteins also requires the assistance of a motor ATPase, SecA, which drives translocation through repeated rounds of ATP binding and hydrolysis (1). In *Escherichia coli,* the catalytic core of SecA contains four domains: nucleotide binding domain-1 (NBD-1; amino acids 1-220 & 378-411), nucleotide binding domain-2 (NBD-2; 412-620), the polypeptide crosslinking domain (PPXD; 221-377) and the α-helical C-terminal domain (CTD; 378-830). SecA binds to substrate proteins in a groove formed between the PPXD and the two NBDs (2). ATP binding and hydrolysis cause conformational changes in the PPXD and CTD that drive protein translocation (3).

In addition to the catalytic core, most SecA proteins contain a relatively long C-terminal tail (CTT). In *E. coli*, the CTT contains a structurally flexible linker domain (FLD; amino acids 831-876 and a small metal-binding domain (MBD; 877-901) (4), which co-purifies with Zn^2+^ (5). The CTT is dispensable for translocation (6-8) although cells producing a truncated version of SecA that lacks the CTT display modest growth and translocation defects (6, 7). The only known function of the MBD (or the CTT generally) is in mediating the interaction of SecA with its binding partner SecB (9, 10), a molecular chaperone that is required for the translocation of a subset of Sec substrates (11). However, the MBD is present in many species that lack a SecB homologue (4, 12), suggesting that it may have an additional function.

Several auxiliary components are also required for efficient Sec-dependent translocation *in vivo*. For example, SecYEG forms a supercomplex with SecD, SecF, YajC and YidC (13, 14), which has been proposed to assist in the insertion and assembly of integral membrane protein complexes (15). Mutations disrupting components of this complex cause broad translocation defects *in vivo* (16-18). In addition, a ribonucleoprotein complex known as the signal recognition particle (SRP) is required for the efficient translocation of a subset of proteins, consisting mostly of integral membrane proteins (19-21). Finally, many soluble periplasmic and outer membrane proteins require the assistance of SecB (22).

Sodium azide inhibits the growth of bacteria by inhibiting SecA (23, 24). Mutations that confer azide resistance map to the *secA* gene (23-25), and azide causes a nearly complete block in SecA-mediated protein translocation within minutes of addition to growing cells (24). Azide inhibits ATP turnover by the F-ATPase by inhibiting exchange of ADP for ATP, and it is thought that azide inhibits SecA by a similar mechanism (26). In support of this notion, most of the mutations that confer azide resistance result in amino acid substitutions in one of the two nucleotide binding domains of SecA (23), and many cause an increased rate of nucleotide exchange *in vitro* (27). However, the concentration of azide required to partially inhibit the rate of ATP turnover by SecA *in vitro* (10-20 mM) (24, 28) is an order of magnitude greater than that required completely block translocation *in vivo* (1-2 mM), suggesting that the mechanism by which azide inhibits SecA is not fully understood.

In this study, we investigated the mechanism of action of azide *in vivo*. Previous genetics studies have relied on either (i) selections for azide resistant mutants or (ii) the characterisation of unrelated mutations in the *secA* gene on azide sensitivity (23, 24, 29, 30). These studies have generally ignored weak effects and effects not attributable to mutations in the *secA* gene. In order to identify these classes of mutation, we screened a very high-density library of *E. coli* transposon insertion mutants (∼1 million independent insertions) for mutants that became enriched or depleted during exponential growth in the presence of azide (31). Consistent with the idea that azide inhibits Sec-dependent protein translocation, insertions in genes encoding many Sec components were depleted during growth in subinhibitory concentrations of azide. In contrast, insertions truncating the CTT of SecA caused *E. coli* to grow faster than the parent strain during exponential growth in the presence of azide. Further investigation indicated that: (i) the CTT of SecA binds to iron; (ii) the CTT is required for high-affinity binding to phospholipids *in vivo*; and (iii) azide disrupts binding of SecA to both iron and phospholipids at physiological concentrations. Our results suggest that azide inhibits SecA by causing it to backtrack in the ATPase cycle, which disrupts its interaction with the membrane.

## Results

### Identification of genes that affect the susceptibility of *E. coli* to sodium azide using TraDIS

We used <u>tra</u>nsposon-<u>d</u>irected insertion-site <u>s</u>equencing (TraDIS) to identify mutations that increased or decreased the susceptibility of *E. coli* BW25113 to azide (32). To this end, we grew a library of ∼1 million independent transposon insertion mutants in the presence of subinhibitory concentrations of azide. Mutations that increase susceptibility to azide became depleted during growth in azide, while mutations that decrease susceptibility to azide became enriched. We then determined the relative number of progeny produced by each insertion mutant by sequencing the transposon junction using Illumina sequencing. Initial experiments indicated that 0.5 mM sodium azide partially inhibited growth while concentrations of 0.25 mM or less had a minimal effect on growth (**supplemental figure S1A**). Consistent with the known activity of azide on protein translocation (24), growth of the TraDIS library in the presence of 0.25 mM or 0.5 mM azide resulted in depletion of mutants containing insertions affecting the Sec machinery (*e.g. yajC*, *secG*, *secF*, *secM*, *yidC*) and cell envelope stress responses (*e.g. cpxR*, *dsbA*) from the library (**figure 1A; supplemental data S1**). However, cells containing insertions in the *secA* and *secB* genes became enriched during growth in azide (**figure 1A**), suggesting that these insertions decreased the susceptibility of *E. coli* to azide. All of the insertions in *secA* resulted in truncation of the CTT of SecA, consistent with the essentiality of the catalytic core for viability. The majority of the insertions enriched during growth in the presence of azide resulted in truncation of SecA between amino acids 822 and 829 at the junction of the CTD and the CTT (**figure 1B**), suggesting that the CTT contributed to azide susceptibility. All of the transposons in *secB* were inserted in the same orientation and maintained expression of the downstream *gpsA* gene (**supplemental figure S1B**) (31). Because increased production of GpsA can suppress some Sec defects (33), we did not investigate the *secB* insertion mutants further. In addition, insertions in genes required for the response to oxidative stress (*e.g. trxC*, *ahpC*, *gor*) were depleted from the library during growth in the presence of azide (**figure 1A; supplemental data S1**), suggesting that long-term exposure to azide causes oxidative stress.

**Figure 1.**
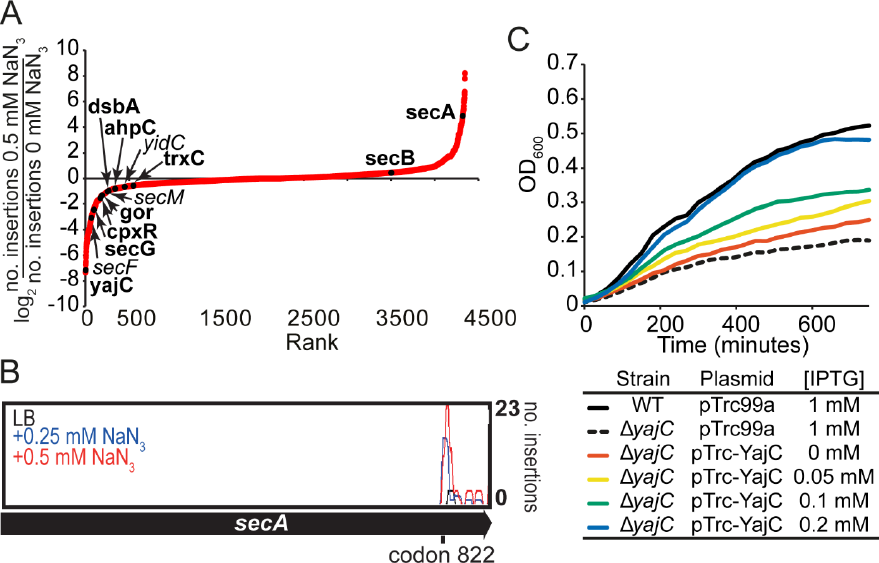
Effect of azide on growth of transposon insertions mutants in TraDIS. (A) Plot of the degree of depletion or enrichment in transposon insertions in all of the non-essential genes in *E. coli* after growth of the TraDIS library to OD_600_ = 0.9 in the presence of 0.5 mM sodium azide (see **supplemental data S1B**). The log_2_ of the fraction of the number insertions in a gene after growth in the presence of 0.5 mM azide over the number of insertions after growth in LB was plotted according to the degree of enrichment. Data points representing insertions in the *secA*, *secB*, *secF*, *secG*, *secM*, *yidC*, *yajC*, *cpxR*, *dsbA*, *ahpC*, *gor* and *trxC* genes are indicated. The growth of selected single gene deletion mutants in the presence of azide (bolded) was compared with that of the parent (see **Table 1**). (B) Number of mutants containing insertions at the indicated location in the *secA* gene after growth of the TraDIS library in the absence (black) or presence of 0.25 (blue) or 0.5 mM (red) NaN_3_. Most of these insertions truncate the *secA* gene between codons 822 and 829 at the junction between the HSD and the CTT. (C) Strains BW25113 (WT) or JW0397 (Δ*yecA*) containing plasmid pTrc99a or pTrc-YajC were grown in the presence of 0.5 mM azide and the indicated concentration of sodium azide in LB at 37^°^C. Growth of the strains was monitored by measuring the increase in optical density at 600 nm.

### Deletion mutations of genes identified in TraDIS have a similar effect on azide sensitivity

To verify the results of our screen, we examined the growth rates of Δ*secG*, Δ*yajC*, Δ*dsbA*, Δ*cpxR*, Δ*gor*, Δ*ahpC* and Δ*trxC* deletion mutants relative to the parent in the presence of azide. With the exception of the Δ*gor* mutant, the presence of 0.25 or 0.5 mM sodium azide had a greater effect on the growth rate of the mutants than it did on the parent strain (**table 1**). Because mutations in *yajC* have not previously been shown to cause defects in protein translocation, we wished to confirm that the increased azide sensitivity of the Δ*yajC* mutant was not caused by polar effects on the expression of the downstream *secD* and *secF* genes (34). IPTG-inducible production of YajC from a plasmid restored growth of the Δ*yajC* mutant to that of the parent in the presence of 0.25 mM sodium azide (**figure 1C**). These results confirmed that it was the lack of YajC in this mutant (and not SecD or SecF) that caused the increased sensitivity of the Δ*yajC* mutant to azide.

**Table 1.** Relative growth rates of single gene deletion mutants from the Keio collection.

| Strain | Growth rates* |  |  |
| --- | --- | --- | --- |
|  | LB<sup>†</sup> | 0.25 mM NaN<sub>3</sub><sup>‡</sup> | 0.5 mM NaN<sub>3</sub><sup>‡</sup> |
| BW25113 | 1.00 ± 0.00 | 1.00 ± 0.09 | 1.00 ± 0.09 |
| *ΔyajC* | 0.88 ± 0.05 | 0.59 ± 0.08 | 0.54 ± 0.11 |
| *ΔsecG* | 1.02 ± 0.05 | 0.53 ± 0.11 | 0.75 ± 0.1 |
| *ΔdsbA* | 1.01 ± 0.08 | 0.95 ± 0.09 | 0.63 ± 0.15 |
| *ΔcpxR* | 1.13 ± 0.13 | 0.93 ± 0.02 | 0.65 ± 0.15 |
| *Δgor* | 0.96 ± 0.01 | 0.97 ± 0.08 | 1.02 ± 0.04 |
| *ΔahpC* | 0.95 ± 0.00 | 0.80 ± 0.06 | 0.64 ± 0.06 |
| *ΔtrxC* | 0.92 ± 0.00 | 0.89 ± 0.02 | 0.78 ± 0.04 |
| MC4100 | 1.00 ± 0.02 | ND<sup>‡</sup> | 1.00 ± 0.03 |
| *secA827::IS1* | 1.03 ± 0.01 | ND<sup>‡</sup> | 1.38 ± 0.18 |

\*Growth rates relative to parent strain grown under the same conditions.
<sup>†</sup>Strains were grown in LB alone or in LB containing the indicated concentration of NaN<sub>3</sub>.
<sup>‡</sup>not determined

We also investigated the effect of azide on the growth of an *E. coli* MC4100 mutant containing an IS*1* insertion at codon ∼827 in *secA* (*secA827*::IS*1*) (6). The *secA827*::IS*1* mutant grew identically to the parent in the absence of azide, but growth of the mutant was significantly less susceptible to the presence of 0.5 mM azide (**table 1**), confirming that truncation of the CTT decreases susceptibility of *E. coli* to azide.

### Azide disrupts binding of the MBD to iron *in vivo*

Structural studies have suggested that the FLD inhibits the interaction of SecA with substrate proteins by binding in the substrate binding groove and that interaction of the MBD with SecB overcomes this inhibition (35, 36). Our results raised the possibility that azide disrupts the structure of the MBD, resulting in auto-inhibition of SecA, which cannot be overcome by the interaction of SecA with SecB. To investigate the effect of azide on the CTT, we purified a protein fusion between the CTT and the small ubiquitin-like modifier (SUMO) *Saccharomyces cerevisiae* (SUMO-CTT) from cells treated with 2 mM sodium azide for 10 minutes (*i.e.* conditions routinely used to inhibit SecA *in vivo*). SUMO-CTT purified from azide-treated cells was more prone to aggregation compared to protein purified from untreated cells (**figure 2A**), suggesting that azide disrupted the structure of the CTT. Because the FLD is structurally flexible, we reasoned that the increased propensity of the CTT to aggregate was likely due to the effect of azide on the structure of the MBD. Because disrupting the structure of the MBD should result in release of the bound metal ion, we determined the metal ion content of SUMO-CTT purified from azide-treated cells using mass spectrometry (ICP-MS) (**figure 2B**). SUMO-CTT copurified with significant amounts of two biologically relevant transition metals: iron and zinc. The samples also contained trace amounts of copper, which was likely a contaminant since copper is not a physiological ligand for cytoplasmic proteins (37). Azide treatment did not significantly affect the amount of zinc that copurified with SUMO-CTT. However, it did cause a strong reduction in the amount of iron that copurified with SUMO-CTT (**figure 2B**). These data suggested that the physiological ligand of the MBD was iron.

**Figure 2.**
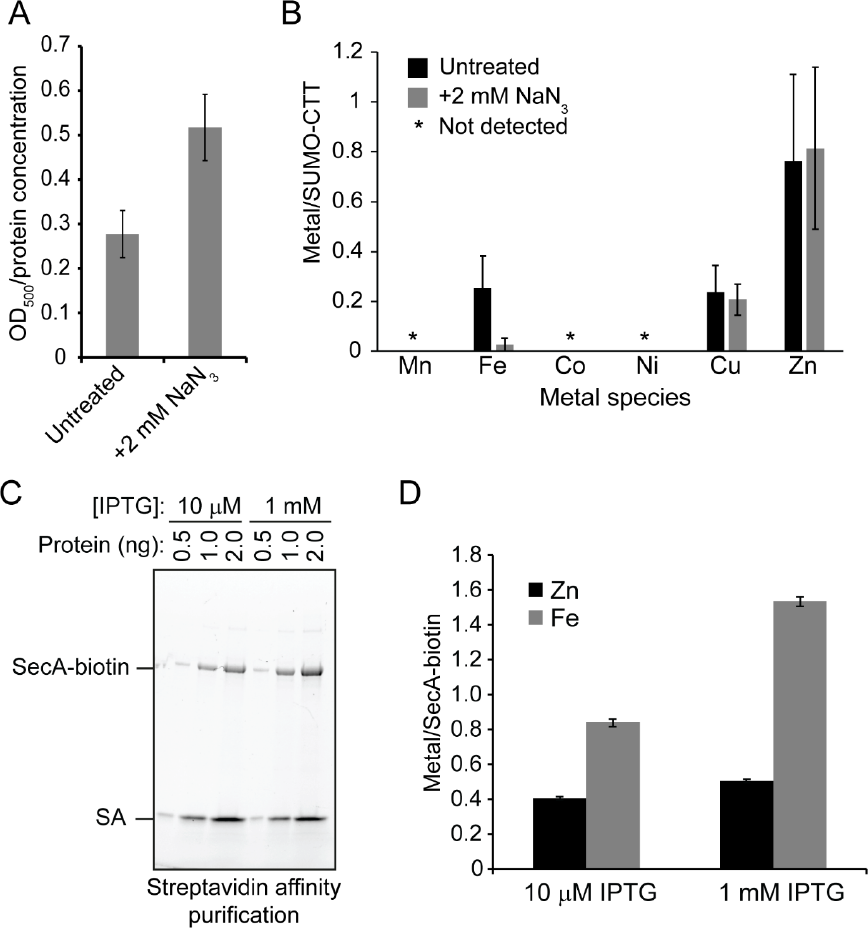
Interaction of SecA with iron *in vivo*. (A & B) Cells producing SUMO-CTT were incubated in the absence (untreated) or presence (+N_3_) of 2 mM NaN_3_ for 10 minutes. SUMO-CTT was purified from the cell lysates using streptactin beads and washed extensively with buffer. (A) The bound protein was eluted from the streptactin beads using 7M guanidinium and was then diluted into buffer lacking guanidinium. Aggregation of the protein was measured using light scattering at 500 nm. Confidence intervals are the s.e.m. (B) The iron content of SUMO-CTT was determined using mass spectrometry (ICP-MS) and normalised to the protein concentration in the eluted protein. Confidence intervals are one s.d. (C & D) Strain DRH839 (*ptrc-secA-biotin*) was grown to OD_600_ = 1 in the presence of either 10 μM or 1 mM IPTG, as indicated. SecA-biotin was purified from the cell lysates using Streptavidin-coupled beads. (C) The eluted protein was resolved by SDS-PAGE on a 15% gel and visualised using BioRad Stain-Free dye. (D) The iron and zinc content of the samples was determined using ICP-OES and normalised to the concentration of protein in the eluted sample. Black, Zn content; Grey, Fe content. Confidence intervals are one s.d.

### SecA copurifies with iron

Because previous studies have suggested that the MBD binds zinc (5, 38), we wished to confirm that full-length SecA binds to iron. Mis-metallation of iron-binding proteins with zinc frequently occurs for several reasons (39, 40): first, the form of iron typically bound by cytoplasmic proteins, Fe^2+^, can be oxidised by dissolved oxygen in purification buffers, while zinc is stable under aerobic conditions; in addition, many iron-metalloproteins have a higher intrinsic affinity for zinc than for iron (37); finally, purification buffers are often contaminated with low (but non-negligible) concentrations of Zn^2+^. To minimize the effect of these issues, we purified SecA as rapidly as possible using a protein that was covalently modified at its C-terminus with biotin (SecA-biotin; (41)). Production of SecA-biotin in strain DRH839 was controlled by an IPTG-inducible promoter, and SecA-biotin was the only version of SecA produced in these cells. Incubation of cells lysates with streptavidin-coupled beads resulted in the purification of only two proteins: SecA-biotin and streptavidin (**figure 2C**). Analysis of the zinc and iron content of the eluted protein using optical emission spectroscopy (ICP-OES) indicated that iron was the dominant metal species present at two different induction levels (10 μM and 1 mM IPTG) (**figure 2D**). These data confirmed that the physiological ligand of the MBD is iron.

### Effect of azide on the oxidation state of the MBD

The MBD coordinates the bound iron *via* three cysteines. Because our genetic analysis suggested that azide causes oxidative stress, we reasoned that that azide might oxidise of the metal-coordinating cysteines, resulting in misfolding of the MBD and release of the bound metal. To investigate this possibility, we examined the effect of azide on binding of the MBD to metal *in vitro*. Because oxidation of iron by oxygen could affect the affinity of SecA for the bound metal, we used full-length SecA that had been charged with Zn^2+^ and measured the rate of release of Zn^2+^ using 4-(2-pyridylazo)resorcinol (PAR). Binding of PAR to Zn^2+^ results in a strong increase in absorbance at 492 nm. The presence of a strong oxidant, hydrogen peroxide, moderately increased the rate of Zn^2+^ release (**figure 3**). However, the presence of 20 mM azide did not detectably affect the rate of Zn^2+^ release by SecA, suggesting that azide does not directly oxidise the metal-coordinating cysteines.

**Figure 3.**
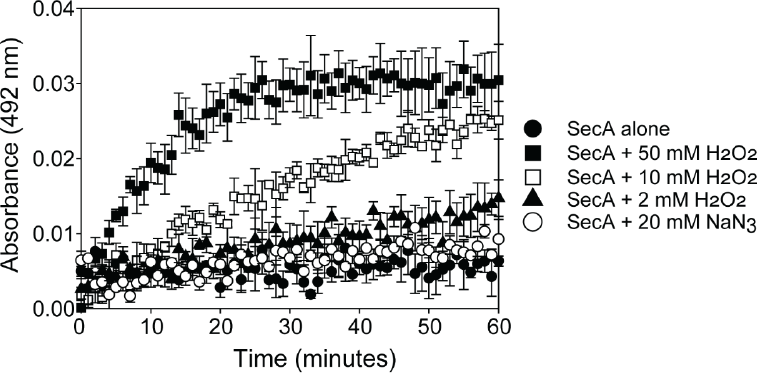
Effect of hydrogen peroxide and sodium azide on metal binding by SecA. 4 μM zinc-bound SecA and 100 μM PAR were incubated in the presence of 2 mM (filled triangles), 10 mM (open squares) or 50 mM (filled squares) hydrogen peroxide, 20 mM sodium azide (open circles) or in the absence of hydrogen peroxide and sodium azide (closed circles). Binding of PAR to free Zn^2+^ was measured by the increase in absorbance at 492 nm over a period of 60 minutes. Confidence intervals are one s.d.

### Azide disrupts binding of SecA to membrane phospholipids

Because the CTT has previously been implicated in binding of SecA to anionic phospholipids (10), we wished to investigate whether azide also disrupted binding of SecA to membrane phospholipids. To this end, we used a hexahistidine-tagged protein fusion between SUMO and SecA (His-SUMO-SecA). This fusion protein was functional *in vivo* since production of the protein from a plasmid could complement the growth defect caused by depletion of SecA (**supplemental figure S2**). His-SUMO-SecA copurified strongly with membrane phospholipids even after several high-salt washes (**figure 4A**), and the copurifying lipids appeared to be enriched for phosphatidylglycerol (PG), consistent with the affinity of SecA for acidic phospholipids (42, 43). In contrast, His-SUMO-SecA containing alanine substitutions in two of the metal-coordinating cysteines (C885 and C887; His-SUMO-SecA^CC/AA^) did not copurify with phospholipids (**supplemental figure S3**), indicating that the MBD is required for high affinity binding to phospholipids. Because our results indicated that azide disrupts the structure of the MBD, we investigated the effect of azide on binding of His-SUMO-SecA to phospholipids. Treating cells producing His-SUMO-SecA with azide prior to lysis strongly disrupted binding of SecA to phospholipids (**figure 4A**).

**Figure 4.**
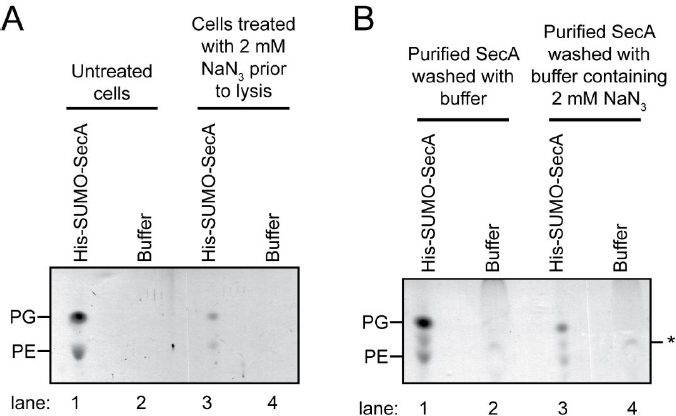
Effect of azide on binding of SecA to phospholipids *in vitro* and *in vivo*. (A) A culture of *E. coli* producing His-SUMO-SecA was grown to late exponential phase. Half of the culture was treated with 2 mM sodium azide for 10 minutes prior to lysis (lanes 3 & 4), and the other half was left untreated (lanes 1 & 2). His-SUMO-SecA was purified from the cell lysates using Ni-affinity purification. Phospholipids from 2 mg of the purified protein were extracted into 100 μl chloroform, and 5μl of the extracted phospholipids (lanes 1 & 3) and the wash buffer (lanes 2 & 4) were resolved using thin-layer chromatography (TLC). The positions of phosphatidylglycerol (PG) and phosphatidylethanolamine (PE) are indicated. (B) His-SUMO-SecA was purified from untreated cells as in (A) and bound to a Ni-NTA beads. The beads were then washed with buffer containing (lanes 3 & 4) or lacking (lanes 1 & 2) 2 mM sodium azide. The lipids from 0.2 mg of protein were extracted from the eluted protein into 100 μl chloroform, and 20 μl of the extracted lipid (lanes 1 & 3) and the wash buffer (lanes 2 & 4) were resolved using TLC. The positions of PG, PE and a non-specific band from the wash buffer (*) are indicated.

To investigate whether azide could disrupt binding of SecA to membranes at similar concentrations *in vitro*, we examined the effect of azide on phospholipid-bound His-SUMO-SecA, which was immobilised on a Ni-NTA column. Washing His-SUMO-SecA with buffer containing 2 mM sodium azide resulted in a rapid loss of the bound phospholipid compared to washing with buffer without azide (**figure 4B**). These results indicated that azide can disrupt binding of SecA to the membrane at physiologically relevant concentrations. Furthermore, because ATP was not added to the wash buffers, the defect in membrane binding was not caused by the inhibition of nucleotide exchange.

## Discussion

Our results indicate that azide does not disrupt SecA-mediated translocation by inhibiting the rate of ATP turnover *in vivo*. Rather, they suggest that azide causes SecA to backtrack from the ADP to the ATP-bound conformation, which disrupts the structure of the MBD and binding of SecA to membrane phospholipids. Normally, SecA cycles through three nucleotide-dependent conformations (**figure 5**): ATP-bound (**yellow**), ADP-bound (**red**) and nucleotide free (**green**). Azide inhibits the F-ATPase by binding to the residues that coordinate the γ-phosphate in the ADP-bound form of the protein, stabilising a conformation that resembles the ATP-bound form, and it has been suggested that azide acts on other ATPases by a similar mechanism (26). The ATP-bound form of SecA has a lower affinity for phospholipids than the ADP-bound form (43), and backtracking to this conformation would result in release of SecA from the membrane. Because binding of SecA to phospholipids is required for protein translocation (42-46), disrupting the interaction of SecA with the membrane would also inhibit protein translocation. If this model is correct, mutations that promote forward progress through the ATPase cycle would confer resistance to azide, which could explain why (i) the vast majority of mutations that confer azide resistance alter residues in NBD-1 and NBD-2 (23) and (ii) so many of these alterations affect the affinity of SecA for nucleotide (27).

**Figure 5.**
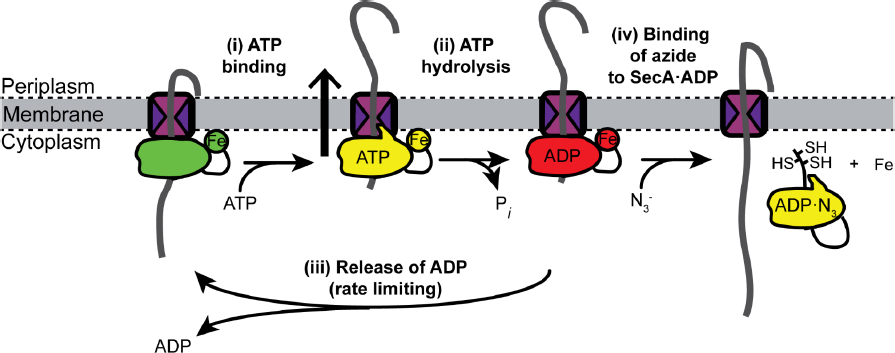
Diagrammatic representation of the mechanism of inhibition of SecA by azide. SecA cycles through three nucleotide bound states: an ATP-bound state (yellow), an ADP-bound state (red) and a nucleotide-free state (green). Binding of SecA to ATP results in translocation of the substrate polypeptide through SecYEG (i) followed by hydrolysis ATP to ADP. Exchange of ADP for ATP requires release of ADP (iii) and is the rate-limiting step the ATPase cycle of SecA. Our results suggest that binding of azide to the ADP-bound form of SecA causes SecA to backtrack to the ATP-bound conformation (yellow), which disrupts binding of SecA to the membrane and to iron (Fe; iv).

The decreased susceptibility of the *secA827* mutant to azide suggests that CTT auto-inhibits the activity of SecA. This result is consistent with previous studies indicating that disruption of the CTT inhibits both binding of SecA to membrane vesicles and translocation of proOmpA *in vitro* (47). Structural studies have suggested that the CTT blocks the interaction of SecA with substrate proteins *in vitro* and that binding of the MBD to SecB relieves this auto-inhibition (35). Disrupting the structure of the MBD would simultaneously release SecA from the membrane and inhibit its interaction with substrate protein. However, because the FLD is required for auto-inhibition (35) but not for translocation (6-8), cells producing a SecA protein lacking the CTT would be less susceptible to azide. It is unclear how azide disrupts the structure of the MBD. However, it is unlikely that azide competes with SecA for binding to iron since *secA827*::IS*1* mutants are only moderately less susceptible to azide than the parent. We speculate that because coordination of the metal ligand by the MBD is strained (4), binding of the MBD to the cytoplasmic membrane stabilises the structure of the MBD. As a result, disrupting the interaction of SecA with the membrane destabilises the structure of the MBD, resulting in release of the bound metal.

Our results strongly suggest that the physiological ligand of the MBD is iron. Previous studies demonstrating that SecA binds to zinc have relied on extensive purification of the protein under aerobic conditions (5, 38). Under these conditions, reaction of the bound iron with oxygen likely destabilises its interaction with the MBD. In addition, because binding of SecA to the membrane could stabilise binding of the MBD to iron, extensive purification of SecA away from phospholipids could lead to loss of the native ligand. Purification buffers are typically contaminated with non-negligible concentrations of zinc and other aerobically stable heavy metals. Indeed, copurification of SUMO-CTT with copper confirms that this sort of exchange takes place during purification. Although copper is equally as effective as zinc at promoting the interaction of the MBD with SecB in *in vitro* binding assays (5), it is not a physiological ligand for cytoplasmic proteins (37).

The increased sensitivity of *yajC* mutants to azide provides the first evidence linking YajC to Sec-dependent protein translocation *in vivo*. Previous studies have attributed defects in translocation caused by mutations in *yajC* to polar effects on expression of *secD* and *secF* (34), which are transcribed in the same poly-cistronic message. However, the ability of a plasmid-born copy of the *yajC* gene to complement a chromosomal deletion of *yajC* indicates that increased azide sensitivity of the Δ*yajC* mutant is caused by a defect in YajC. This phenotype provides a new tool for investigating the function of YajC *in vivo*. These results demonstrate the power of TraDIS for identifying hidden or subtle phenotypes.

## Methods

### Chemicals and media

All chemicals were purchased from Fisher or Sigma-Aldrich unless indicated. Cells were grown using LB (48). Where noted, sodium azide and isopropylthiogalactoside (IPTG) were added at the indicated concentration. Where required, ampicillin (200 μg/ml) or kanamycin (30 μg/ml) were added to the growth media.

### Strains and plasmids

A description of all the strains and plasmids used in this study can be found in **supplemental table S1**. Strains and plasmids were constructed using common genetic methods (48, 49). Single gene deletion mutants from the Keio collection (50) were obtained from the *E. coli* genetic stock centre (CGSC; Yale University, New Haven, Connecticut). Strains DRH745 and DRH839 were constructed by integrating the IPTG-inducible copy of *secA* on plasmid pDH692 or pDH771 onto the chromosome of MC4100 at the phage λ attachment site using λInCh (51) and transducing the Δ*secA*::Kan mutation into this strain using P1. Strain MG1115 was a kind gift from M. Grabowicz and T. Silhavy. Plasmid pDH543 was constructed by PCR amplifying a fragment of the *secA* gene encoding the C-terminal 70 amino acids and cloning into plasmid pCA597 cut with *BsaI* and *BamHI* (52). Plasmid pTrc-YajC was constructed by cloning a PCR-amplified DNA fragment corresponding the *yajC* coding sequence into strain pTrc99a (Promega).

### TraDIS

TraDIS was carried out similar to Langridge*, et al.* (32). 50 ml of LB broth containing 0, 0.25 or 0.5 mM NaN3 were inoculated with 10 μl of a library of ∼1 million *E. coli* BW25113 mini-Tn*5* insertion mutants, and the cultures were grown to OD_600_ 1.0. Genomic DNA was extracted using a Qiagen QIAamp DNA blood mini kit and then processed using a two-step PCR method (53), which results in Illumina-compatible products. The PCR products were purified using the Agencourt AMPure XP system by Beckman Coulter. The products were sequenced using an Illumina MiSeq sequencer and the reads were mapped to the *E. coli* reference genome NC_007779.1 (*E. coli* K-12 substr. W3110). The number of insertions in the coding sequences (CDS) for each gene was then determined. To reduce the number of false positives due to sequencing assignment errors, genes that exist in multiple copies on the chromosome (e.g. insertion elements and rRNA operons) were eliminated from the analysis. In addition, genes containing 15 or fewer total insertions across all three conditions were eliminated in the data presented in **figure 1A**. BED files of the aligned sequences, the sequence file used for alignments and the features file used for assessing the location of the insertions are available at figshare.com (doi: 10.6084/m9.figshare.5280733).

### Determination of metal content of SecA-biotin and Strep-SUMO-CTT

To determine the effect of azide on binding of Strep-SUMO-CTT to iron, 100 ml cultures of BL21(DE3) containing plasmid pDH543 was grown to OD_600_ 1.0 in LB, and production of Strep-SUMO-CTE was induced using 1 mM IPTG. After 1 hour, cultures were split, and half the culture was treated with 2 mM NaN_3_ for 10 minutes. Cells were rapidly cooled, harvested by centrifugation and lysed using B-PER cell lysis reagent from Pierce (Rockford, Illinois). Cell lysates were incubated for 15 minutes with 50 μl of a 50% Streptactin-Sepharose slurry that had been pre-equilibrated with wash buffer (10 mM HEPES (potassium salt) pH 7.5, 100 mM potassium acetate) and washed extensively with wash buffer. The samples were washed a final time using 10 mM HEPES (potassium salt) pH 7.5 to remove excess salt, and the total protein was eluted off of the column using using 10 mM HEPES (potassium salt) pH 7.5 buffer containing 7M guanidinium hydrochloride. The propensity to aggregate was determined by diluting 50 μl of the guanidinium-denatured protein into 950 μl of a 20 mM HEPES buffer and measuring light scattering at 500 nm. The metal content of the samples was determined using inductively coupled plasma-mass spectrometry (ICP-MS) (School of Geography, Earth and Environmental Sciences, University of Birmingham, UK).

To determine the metal content of full-length SecA-biotin, a 100 ml culture of strain DRH839 was grown in the indicated concentration of IPTG to OD_600_ ∼ 1 and lysed using cell disruption. Cell lysates were incubated with 100 μl Streptactin-sepharose for 15 minutes on ice, and the sepharose beads were washed four times with 30 ml binding buffer (50 mM HEPES, pH 7.5, 100 mM K·acetate, 10 mM Mg·acetate, 0.1% nonidet P40). Metal was eluted from the beads by incubating with metal elution buffer (10 mM HEPES, pH 7.5, 50 mM EDTA) at 55^°^C for 30 minutes, and the zinc and iron content were determined using inductively coupled plasmid-optical emission spectroscopy (ICP-OES). Protein was eluted from the beads by boiling in SDS sample buffer, and the protein content was determined using a Bradford assay. The eluted protein was resolved on a BioRad 15% TGX gel, and the protein bands were visualised using the BioRad Strain-Free stain.

### In vitro zinc-release assay

Full-length SecA was purified similar to Huber*, et al.* (54) except that the protein was not subjected to an on-column denaturation step. SecA was charged with Zn^2+^ by incubating 40 μM of the purified protein with an equimolar concentration of ZnCl_2_. Excess metal was removed from the protein using a 5 ml HiTrap Desalting Column (GE Healthcare). To measure the rate of Zn^2+^ release, 4 μM SecA was incubated in buffer containing 20 mM HEPES (potassium salt), pH 7.5, 100 mM potassium acetate, 20 mM magnesium acetate and 100 μM PAR in the presence of the indicated concentration of H_2_O_2_ or NaN_3_. The increase in PAR absorbance was measured every 60 s for 90 minutes at 492 nm using a BMG Labtech FLUOStar plate reader.

### Phospholipid binding assay

Strain DRH625 or MJ118 was grown to late log phase in LB, and production of His-SUMO-SecA was induced using 1 mM IPTG. Where indicated, the culture was divided after the addition of IPTG, and half of the culture was treated with 2 mM sodium azide for 10 minutes. Cells were harvested by centrifugation, resuspended in lysis buffer (20 mM HEPES [potassium salt] pH7.5, 500 mM NaCl, 1 mM tris(2-carboxyethyl)phosphine [TCEP]) and lysed by cell disruption. Clarified cell lysates were passed over a 1 ml Ni-HiTrap column (GE Biosciences), and the bound protein was washed with 50 volumes of lysis buffer containing 20 mM Imidazole. His-SUMO-SecA was eluted from the column using lysis buffer containing 500 mM imidazole and dialysed against wash buffer lacking imidazole. The concentration of the protein was adjusted to 1 mg/ml to a final volume of 2 ml, and the phospholipids were extracted using 2 ml methanol and 1 ml chloroform. The chloroform phase was removed, and phospholipids were concentrated by air drying and resuspending in chloroform to a final volume of 100 μl. The indicated volume of sample was spotted onto TLC silica gel 60 T_254_ plates (Merck Millipore) and resolved using a mixture of 60 chloroform: 25 methanol: 4 water. For the *in vitro* phospholipid release assay, 0.2 mg of His-SUMO-SecA purified from untreated cells was bound to 100 μl Ni-NTA agarose beads (Life Technologies). The beads were incubated with 500 μl of lysis buffer or buffer containing 2 mM sodium azide for 10 minutes and washed three times with 500 μl lysis buffer. The protein was eluted from the beads with lysis buffer containing 500 mM imidazole, and the phosopholipid content of the supernatant was analysed using the method described above.

## Acknowledgements

We thank Dr. S. Baker and Dr. M. Thompson for technical assistance. We also thank Dr. J. Bryant, Dr. T. Knowles, E. Goodall, Prof. J. Cole and members of the Huber, Henderson, Lund and Grainger labs for helpful advice and discussions. This work was funded by the Biotechnology and Biological Sciences Research Council grant BB/L019434/1 to D. H., M. J. and R. C..

---

**Table S1.**
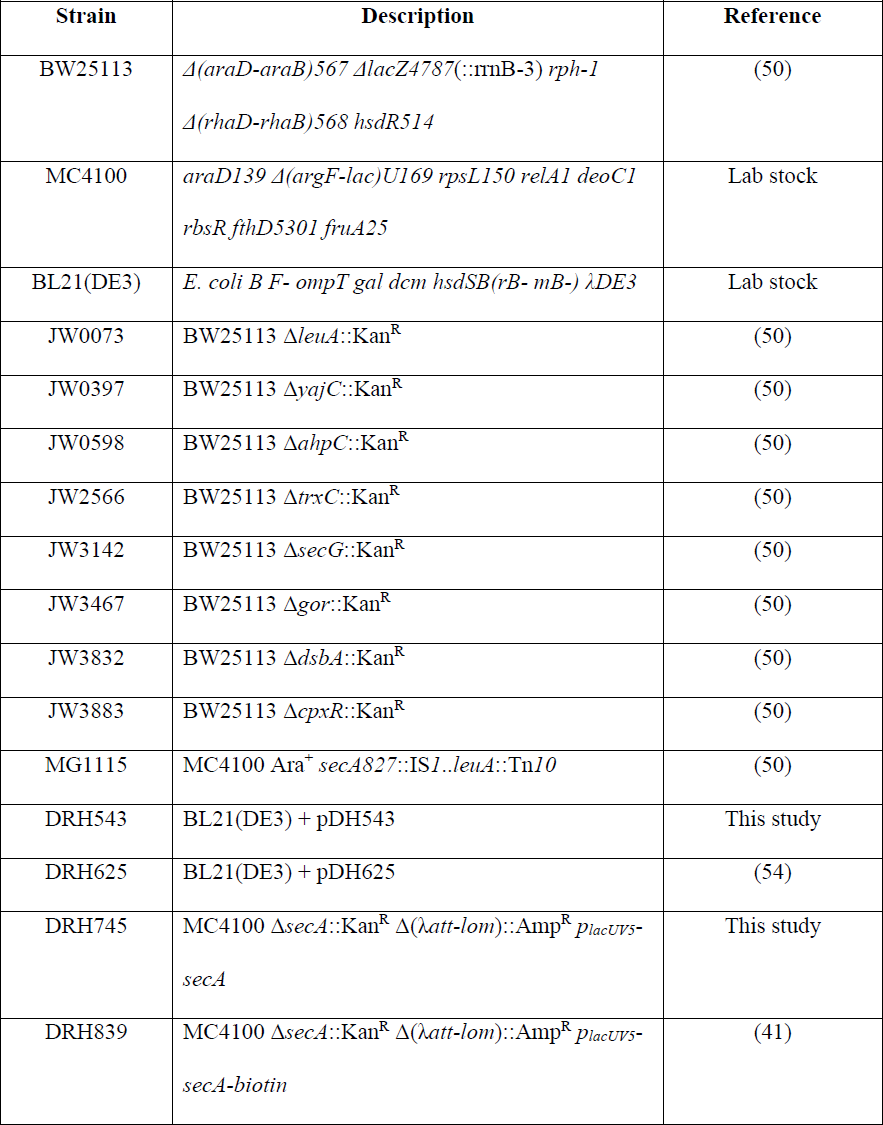
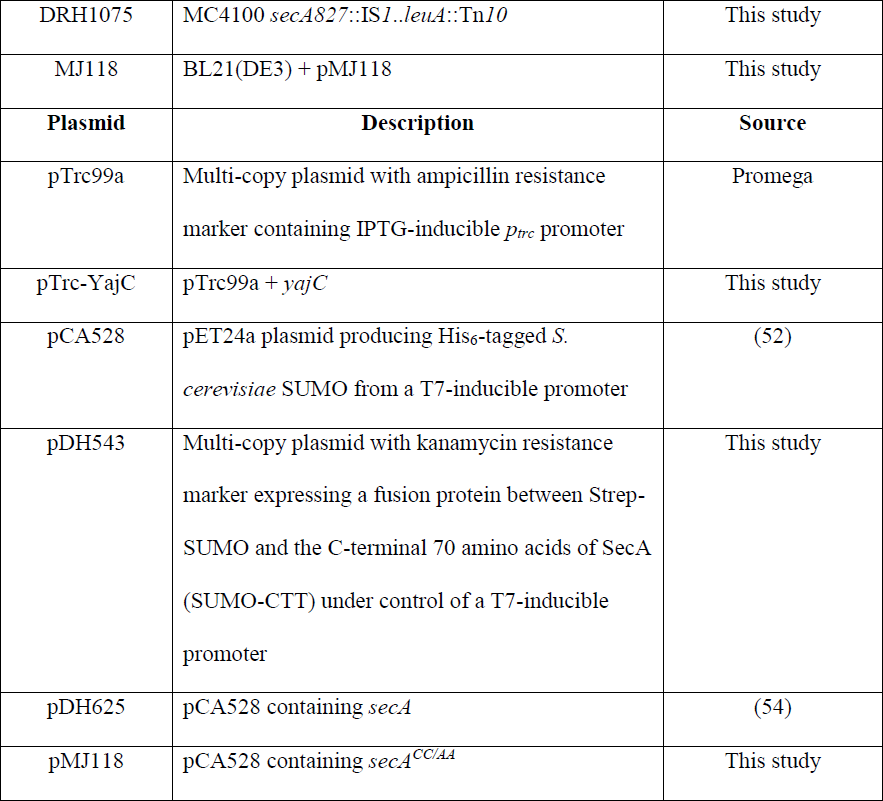
Strains and plasmids used in this study.

**Figure S1.**
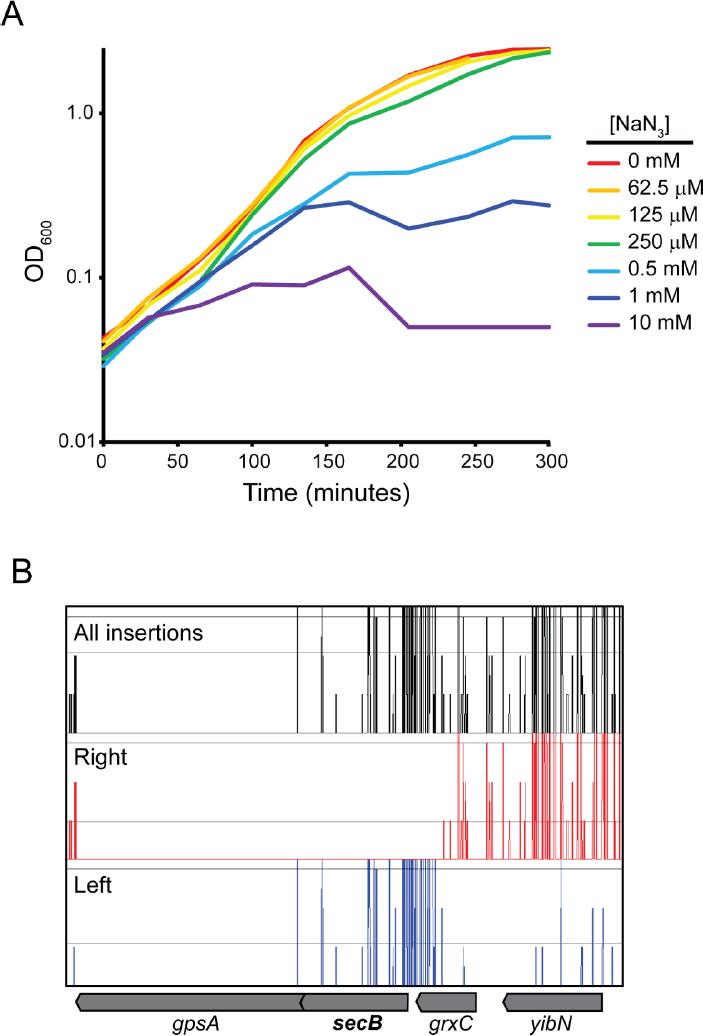
TraDIS analysis of the effect of azide on the growth of *E. coli* BW25113. (A) Strain BW25113 was grown in LB in the presence of the indicated concentration of sodium azide. Growth of the culture was monitored by optical density at 600 nm. (B) Map of unique transposon insertions in the *yibN*-*grxC*-*secB*-*gpsA* operon. Depicted are the location of all transposons in the TraDIS transposon library (black), transposons with the internally encoded promoter transcribing rightwards (red) and transposons with the internally encoded promoter transcribing leftward (blue). The *gpsA* gene contains no insertions suggesting it is essential under these growth conditions. In addition, all of the insertions in the *secB* gene preserve expression of the downstream *gpsA* gene.

**Figure S2.**
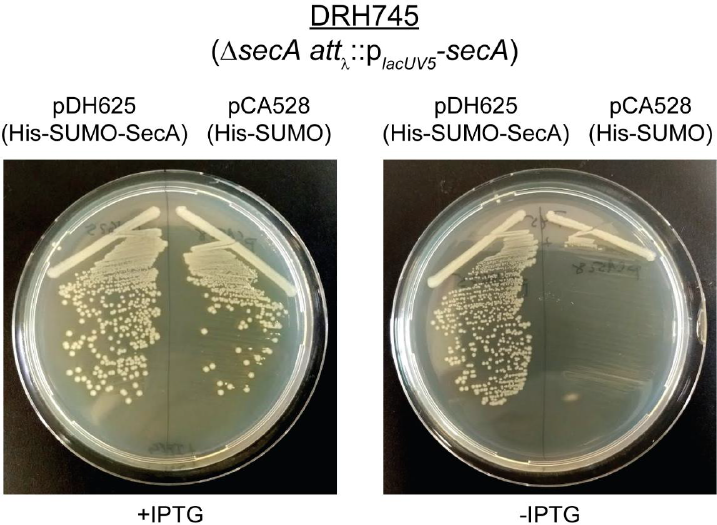
Production of His-SUMO-SecA can complement the growth defect caused by depletion of SecA. Strain DRH745 (p_lacUV5_-*secA*) containing either pDH625 or pCA528 was restreaked onto LB plates containing 1 mM IPTG or lacking IPTG and incubated overnight at 37^°^C.

**Figure S4.**
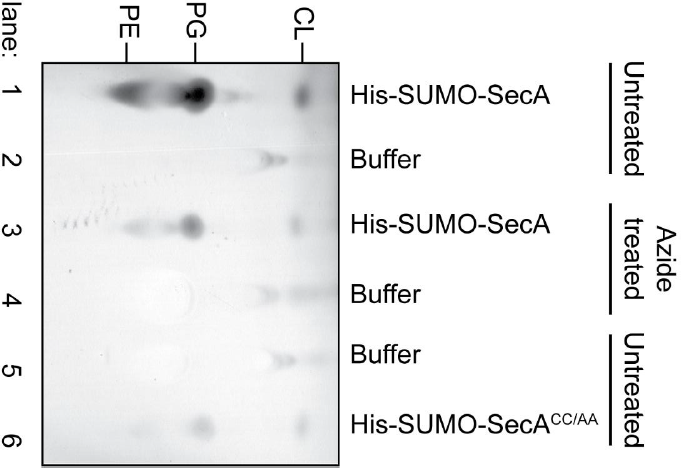
Effect of alanine substitution that disrupt the structure of the MBD on phospholipid binding. Cultures of *E. coli* producing His-SUMO-SecA (lanes 1-4) or His-SUMO-SecA^CC/AA^ (lanes 5& 6) were grown to late exponential phase. In the case of His-SUMO-SecA, half of the culture was treated with 2 mM sodium azide for 10 minutes prior to lysis (lanes 3 & 4), and the other half was left untreated (lanes 1 & 2). His-SUMO-SecA and SUMO-SecA^CC/AA^ were purified from the cell lysates using Ni-affinity purification. Phospholipids from 2 mg of the purified protein were extracted into 100 μl chloroform, and 5μl of the extracted phospholipids (lanes 1, 3 and 6) and the wash buffer (lanes 2, 4 and 5) were resolved using thin-layer chromatography (TLC). The positions of phosphatidylglycerol (PG), phosphatidylethanolamine (PE) and cardiolipin (CL) are indicated.

**Supplemental Data S1. Analysis of transposon insertion mutants in TraDIS library after outgrowth in LB or LB containing NaN_3_.** (A) Number of insertions in each gene in the *E. coli* genome after growth of the TraDIS library in LB, LB containing 0.25 mM NaN_3_ or LB containing 0.5 mM NaN_3_. Genomic DNA from one experiment was prepared and sequenced twice to increase the number of sequencing reads and to examine the reproducibility of the results. For each run the number of insertions in each gene was normalised to the total number of sequencing reads. (B) The degree of enrichment or depletion of insertion mutations in the library after growth in the presence of azide was calculated by determining the log (base 2) of the ratio of the number of insertions in a gene after outgrowth in the azide-treated conditions to the number of insertions after outgrowth in LB. The BED files used to derive the data presented in parts A and B is available at figshare.com (doi: 10.6084/m9.figshare.5280733).

